# HTRX: an R package for learning non-contiguous haplotypes associated with a phenotype

**DOI:** 10.1101/2022.11.29.518395

**Authors:** Yaoling Yang, Daniel Lawson

**Affiliations:** Department of Statistical Science, School of Mathematics, University of Bristol, Bristol, BS8 1UG, United Kingdom; Integrative Epidemiology Unit, Population Health Sciences, University of Bristol, Bristol, BS8 2BN, United Kingdom

## Abstract

**Summary:** Haplotype Trend Regression with eXtra flexibility (HTRX) is an R package which uses cross-validation to learn sets of interacting features for a prediction. HTRX identifies haplotypes composed of non-contiguous single nucleotide polymorphisms (SNPs) associated with a phenotype. To reduce the space and computational complexity when investigating many features, we constrain the search by growing good feature sets using ‘Cumulative HTRX’, and limit the maximum complexity of a feature set.

**Availability:** HTRX is implemented in R and is available under GPL-3 license from CRAN and Github at: https://github.com/YaolingYang/HTRX.

## 1 Introduction

Numerous single nucleotide polymorphisms (SNPs) associated with human complex traits and diseases have been discovered by genome-wide association studies (GWAS) (Buniello *et al*., 2019). However, multiple SNPs from a high linkage disequilibrium (LD) region may be required to describe the total causal signal (Yang *et al*., 2012) which may be from multiple SNPs or interactions (Galarneau *et al*., 2010). Haplotype-based association studies, combining SNPs into features, have the potential to be more powerful than methods based on independent SNPs (Balliu *et al*., 2019). Haplotype-based association studies, such as Haplotype Trend Regression (HTR) (Zaykin *et al*., 2002) (reviewed by Schaid (2004) and Liu *et al*. (2008)) are limited to investigating haplotypes specifying values at all SNPs. We recently (Barrie *et al*., 2022) proposed Haplotype Trend Regression with eXtra flexibility (HTRX) which searches non-contiguous haplotypes. As the number of haplotypes increases exponentially with the number of SNPs, inferring true interactions at scale is unrealistic (Guan and Stephens, 2011). Consequently, the goal of HTRX is to make good predictions by selecting features which have the best predictive performance. By predicting the out-of-sample variance explained (*R*^2^), HTRX quantifies whether a tagging SNP is adequate, or whether interactions or LD with unobserved causal SNPs are present.

Barrie *et al*. (2022) demonstrated the utility of this method by detecting interactions in the human leukocyte antigen (HLA) locus for Multiple Sclerosis (MS). This note addresses two important improvements: controlling computational complexity, and ensuring that overfitting is controlled. We will control the former by limiting the flexibility of haplotypes to be considered, and the latter using penalisation and cross-validation (CV). In addition to Bayesian Information Criteria (BIC) (Schwarz, 1978) employed to select candidate models by Barrie *et al*. (2022), we consider Akaike’s information criterion (AIC) (Akaike, 1974) and lasso (Tibshirani, 1996) regularisation in the R package ‘HTRX’.

## 2 Methods

HTRX defines a template for each haplotype using the combination of ‘0’, ‘1’ and ‘X’ which represent the reference allele, alternative allele and either of the alleles, respectively, at each SNP. For example, a four-SNP haplotype ‘1XX0’ only refers to the interaction between the first and the fourth SNP. Each haplotype *H*_*ij*_ takes a value of either 0, 0.5 or 1 if individual *i* has 0, 1 or 2 haplotype *j* in both genomes. This template creates 3^*u*^ *−* 1 different haplotypes in a region containing *u* SNPs, while only 2^*u*^ *−* 1 of them are independent. Fitting models using all possible haplotypes result in overfitting (Hawkins, 2004). To address this, HTRX considers AIC, BIC and lasso penalisation. AIC and BIC penalise the number of features in the model through forward regression, while lasso uses L1 norm to regularize parameters, and the features whose parameters do not shrink to 0 are retained. The objective function of HTRX is the out-of-sample variance explained by haplotypes within a region. We use *k*-fold CV, an ensemble learning method (*k* ≥ 3 for algorithms below) as a natural score function:

### Algorithm 1

Direct CV (function ‘do direct cv’)

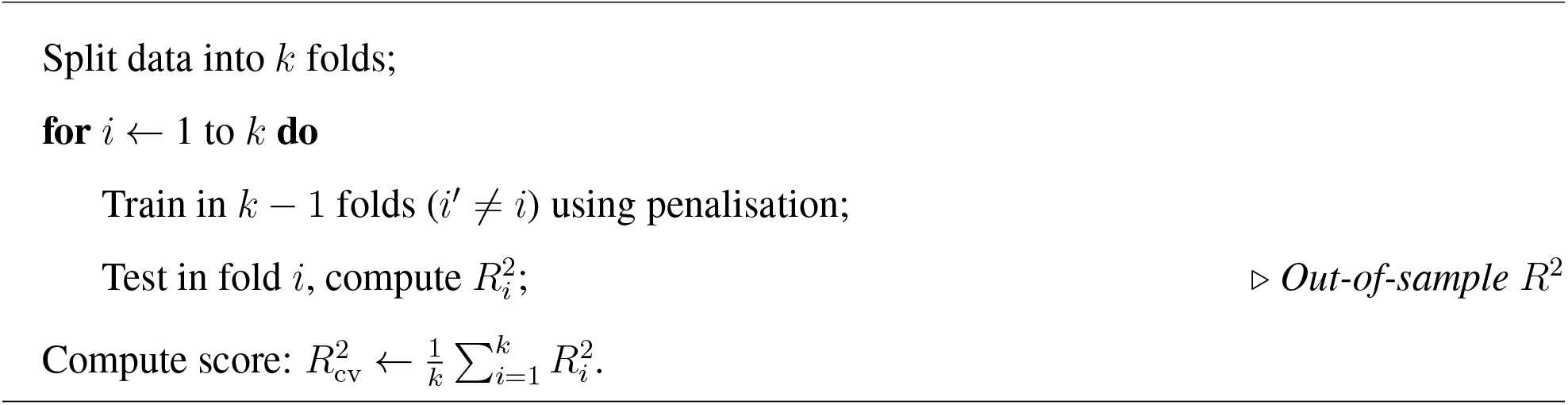

Although Algorithm 1 reduces overfitting, it generates an ensemble of feature sets due to variation in the training data. To choose a fixed feature set, a more complicated CV algorithm is required.

### Algorithm 2

Two-stage CV (function ‘do cv’)

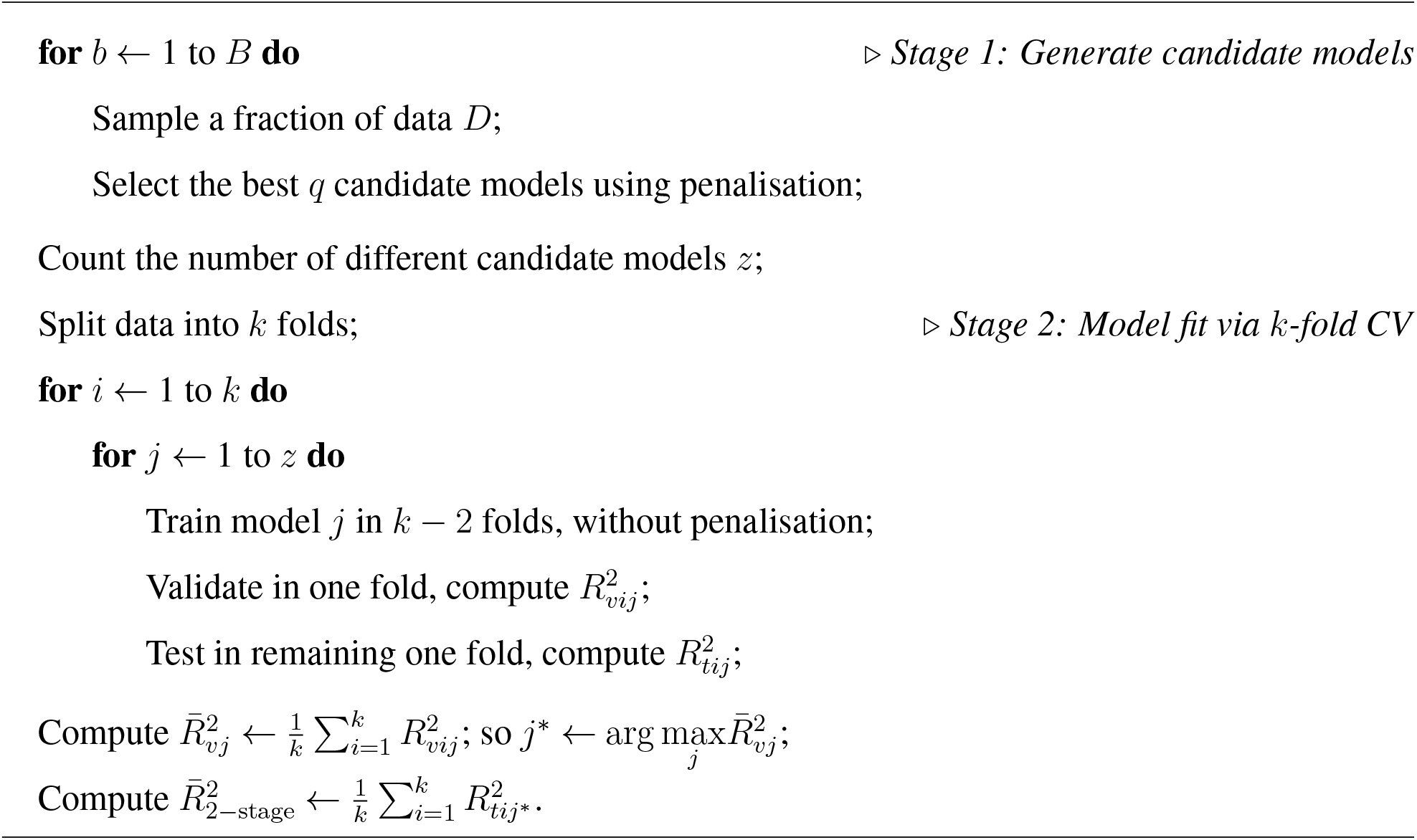

These algorithms scale badly with *u* are limited to around *u ≤* 6. To consider more features as commonly found in genetic LD blocks, *cumulative* HTRX (Algorithm 3) controls the space and computational complexity:

### Algorithm 3

Cumulative HTRX (function ‘do cumulative htrx’)

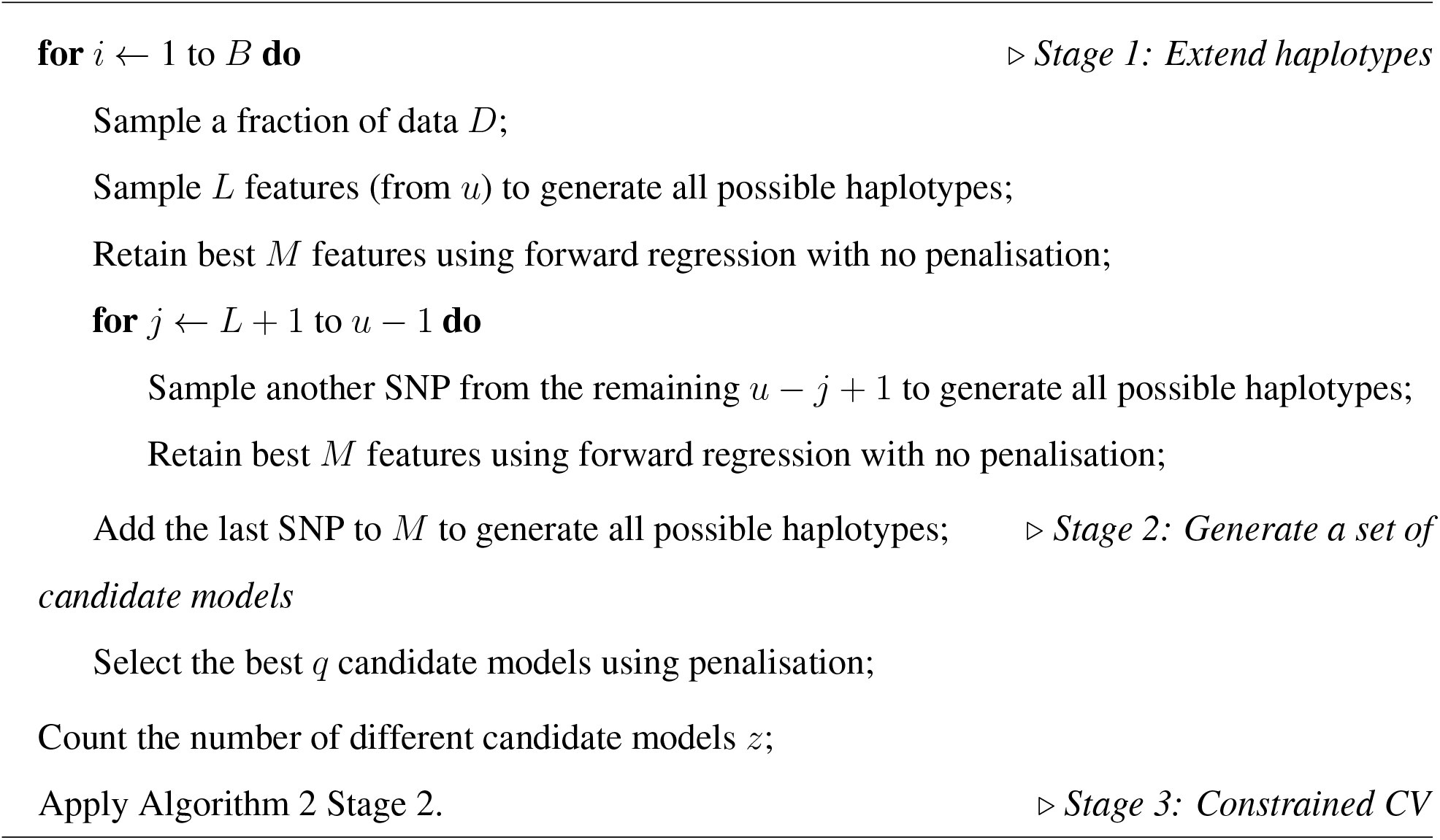

Larger *L, M* and *B* may slightly improve the predictive performance, but they significantly increase complexity both the spatially and computationally. It is rare that many features are involved in an interaction, and such interactions are statistically hard to identify. Because of this, we consider constraining the maximum number of SNPs permitted in a template. This reduces compute, and space complexity (Fig. S1). Fig. 1 compares the performance of many choices in a realistic simulated scenario (Supplementary Methods).

**Figure 1:**
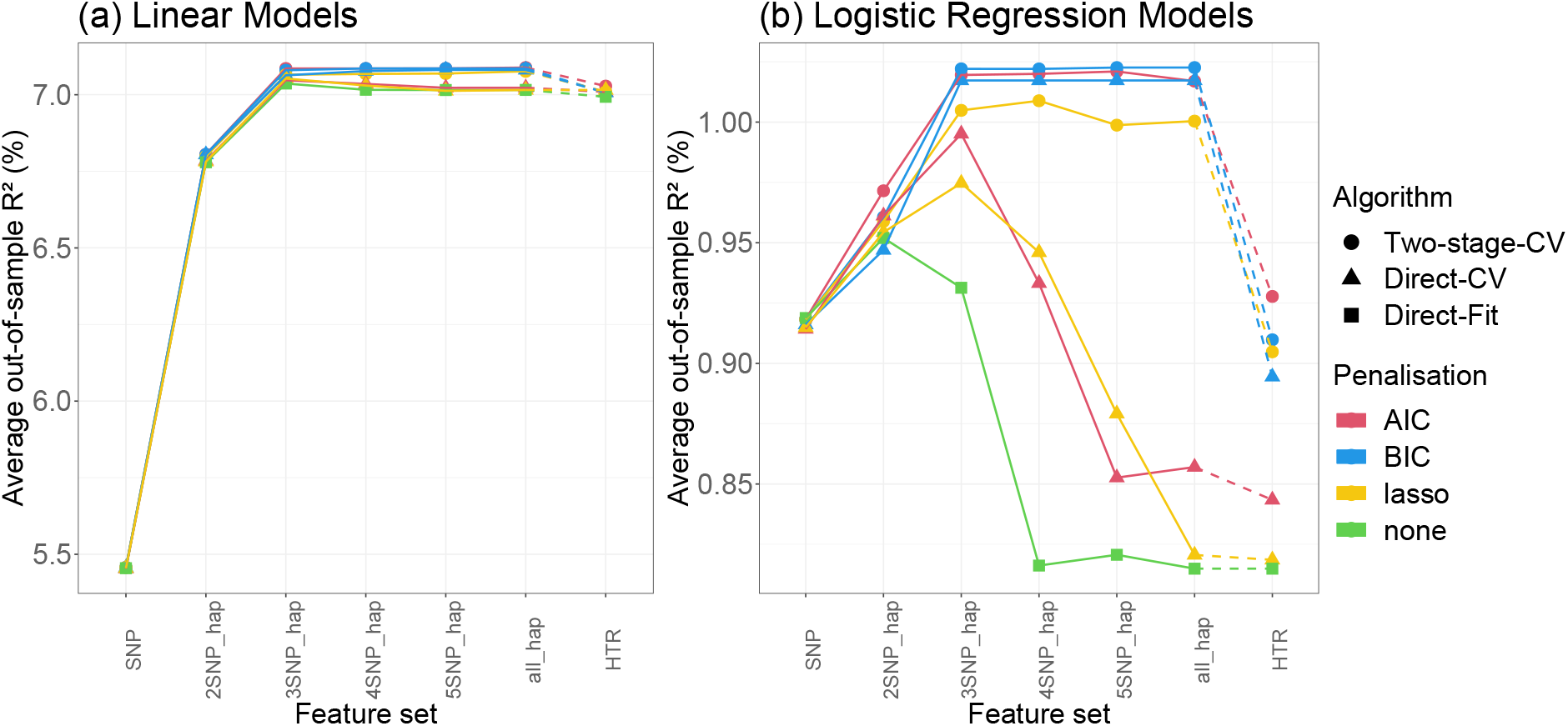
Comparison of the average out-of-sample *R*^2^ through 10-fold CV for linear and logistic regression models in a simulated dataset (Supplementary Methods). Feature set specifies the maximum number of features (of 6) that can interact, from ‘SNP’= 1, to ‘all hap’= 6, while ‘HTR’ uses templates that interact in all SNPs with no ‘X’ in the template. *D* = 50%, *B* = 10 and *q* = 3 are used for ‘Two-stage-CV’. ‘Direct-Fit’ refers to all-feature multivariate regression.

## 3 Discussion

We show in a simulated dataset of 6 interacting features that ‘Two-stage CV’ significantly outperforms ‘Direct CV’, especially when the outcome is binary (Fig. 1). Also, penalisation using AIC or BIC produces significantly better out-of-sample performance than lasso. ‘Two-stage CV’ chooses a subset of all possible models to reduce computational cost.

Out-of-sample prediction is wasteful of scarce data, and for *R*^2^ prediction using less data creates a downward bias as well as increased variance. Whilst results throughout are shown for true out-of-sample prediction, for real data we implement *k*-fold Cross-Validation which introduces a negligible bias compared to the variance (Fig. S2).

More generally, these algorithms efficiently search features for interactions. One approach is reducing the number of interactions permitted, and another is growing the most promising interaction sets. Both exploit the diminishing marginal returns of complexity for prediction to quantify the *total out-of-sample variance explained*, which tests for the presence of feature interaction. It has general application in regression, but is specifically important for determining whether a single SNP is an adequate description of the effect of a genetic region on a phenotype.

