## Supplementary Methods for "HTRX: an R package for learning non-contiguous haplotypes associated with a phenotype"

### 1 Supplementary Methods

#### 1.1 Simulation

We simulate 6 biallelic SNPs $G_{ij}$ ($i = 1, \ldots, 100,000$ denotes individuals and $j = 1, \ldots, 6$ denotes SNPs) with frequency about 20% for the alternative allele '1' from an LD block, i.e. the correlation between each pair of SNPs is around 97.8%. We assumed each individual is haploid, and created all possible haplotypes in the 6-SNP region.

We generate the effect size of two SNPs $G_{\cdot 2}$ and $G_{\cdot 4}$ as $\beta_{G_2} = \frac{0.5}{sd(G_{\cdot 2})}$ and $\beta_{G_4} = \frac{0.5}{sd(G_{\cdot 4})}$. We only select two haplotypes with real effects: a two-SNP interaction $H_{\cdot 1}$ ('X0XX1X') with effect size $\beta_{H_1} = \frac{0.3}{sd(H_{\cdot 1})}$ and a three-SNP interaction $H_{\cdot 2}$ ('1XX0X1') with effect size $\beta_{H_2} = \frac{0.3}{sd(H_{\cdot 2})}$. Besides, we create a confounder $C = 0.5G_{\cdot 2} - 0.8G_{\cdot 4}$ which has effect size $\beta_C = \frac{1}{sd(C)}$. A random error with large variance $e_i \sim N(0, 4)$ and an intercept term $\lambda = -4$ are also generated. We investigate both continuous and binary outcomes using linear and logistic regression models. When the outcome $Y^c$ is continuous,

$$Y_i^c = \beta_{G_2} G_{i2} + \beta_{G_4} G_{i4} + \beta_{H_1} H_{i1} + \beta_{H_2} H_{i2} + \beta_C C_i + \lambda + e_i.$$

For binary outcome, we sample $Y_i^b \sim \text{Bin}(1, \pi_i)$ where $\pi_i = \frac{e^{Y_i^c}}{1+e^{Y_i^c}}$ is the probability that the $i$th individual has the outcome.

Then we compare the out-of-sample performance of different algorithms, penalisation and models with parameters $k = 10$, $D = 50\%$, $B = 10$ and $q = 3$ (Fig. ??). The comparison results of different algorithms, penalisation and models have been included in the main text.

#### 1.2 Algorithm remark

In this subsection, we justify the recommended algorithm 'Two-stage CV' for short regions, which automatically justifies the algorithm 'Cumulative HTRX' for longer regions. In the first stage, we sampled a subset of all possible models as the candidate models, from which we select the best model through $k$-fold cross-validation in the second stage. Generating more candidate models can improve the estimate accuracy, which can be realised by increasing simulation times ($B$) or the number of best models we keep ($q$). Furthermore, We recommend a small percentage of $D$, because sub-data with larger variation increases the possibility that different candidate models are selected.

In the second stage, we split the dataset into $k$ folds ($k \geq 3$), and stratified sampling is applied for binary outcome. We make a train ($k - 2$ folds), validation (1 fold) and test (1 fold) data split for each time of the CV loop. The training data is used for fitting each candidate model, and the out-of-sample $R^2$ is computed on the validation data. Also, we record the out-of-sample $R^2$ computed on the test set. After repeating the process $k$ times when each fold has been used as the validation data once, we select the best model $j^*$ which has the maximum average out-of-sample $R^2$ in the validation data. We finally report the average out-of-sample $R^2$ of each test set, which is independent of the training and validation set in each time of the $k$-fold CV, although it is used for training and validation in the other times. In principle there is a potential for this re-use of data to cause a bias. We don't split an entirely independent test data in the HTRX package, because our goal is to a) estimate out-of-sample $R^2$ in order to b) rank models containing interactions vs not. By averaging over $k$-folds, the cross-validation procedure reusing parts of the entire dataset leads to more accurate (i.e. lower root-mean-square error) estimates.

We illustrate through simulation (Fig. S2) that the within-CV average out-of-sample $R^2$ is a good estimator of the average out-of-sample $R^2$ tested on entirely independent datasets. Using the parameters in Supplementary Methods 1.1 and features selected from all possible haplotypes penalised by either AIC, BIC or lasso through 'Two-stage CV' for linear or logistic regression models, we compare model performance (out-of-sample $R^2$) in the following setups:
(1) Algorithm 2 but testing on only one fold of the CV data, as would be obtained by using a traditional training/test split;
(2) Algorithm 2 using full Cross-Validation, our recommended approach for real data;
(3) Follow Algorithm 2 by training on all the CV data, but test on an independent new dataset with the same size as the CV data, which requires double the data size.

Then we compare the average and standard deviation of the out-of-sample $R^2$ obtained by different setups in Fig. S2. The mean values of each out-of-sample $R^2$ are similar, while testing on a single set of the CV data has a significantly larger standard deviation than the average of $R^2$ tested on each fold of the CV data. The $R^2$ for setup (3) is the model performance on an entirely large independent dataset while trained on all the CV data. As the training data for setup (3) is larger than (1) and (2), the fitted model should be more reliable. Compared with the $R^2$ for setup (3), the within-CV average out-of-sample $R^2$ that we report in 'Two-stage CV' and 'Cumulative HTRX for CV' (the $R^2$ for setup (2)) has almost equivalent distribution for linear outcomes and minor underestimation for binary outcomes. Therefore, our estimated average out-of-sample $R^2$ exhibits little, if any, bias while avoiding power loss by splitting entirely independent test data.

#### 2 Supplementary Figures

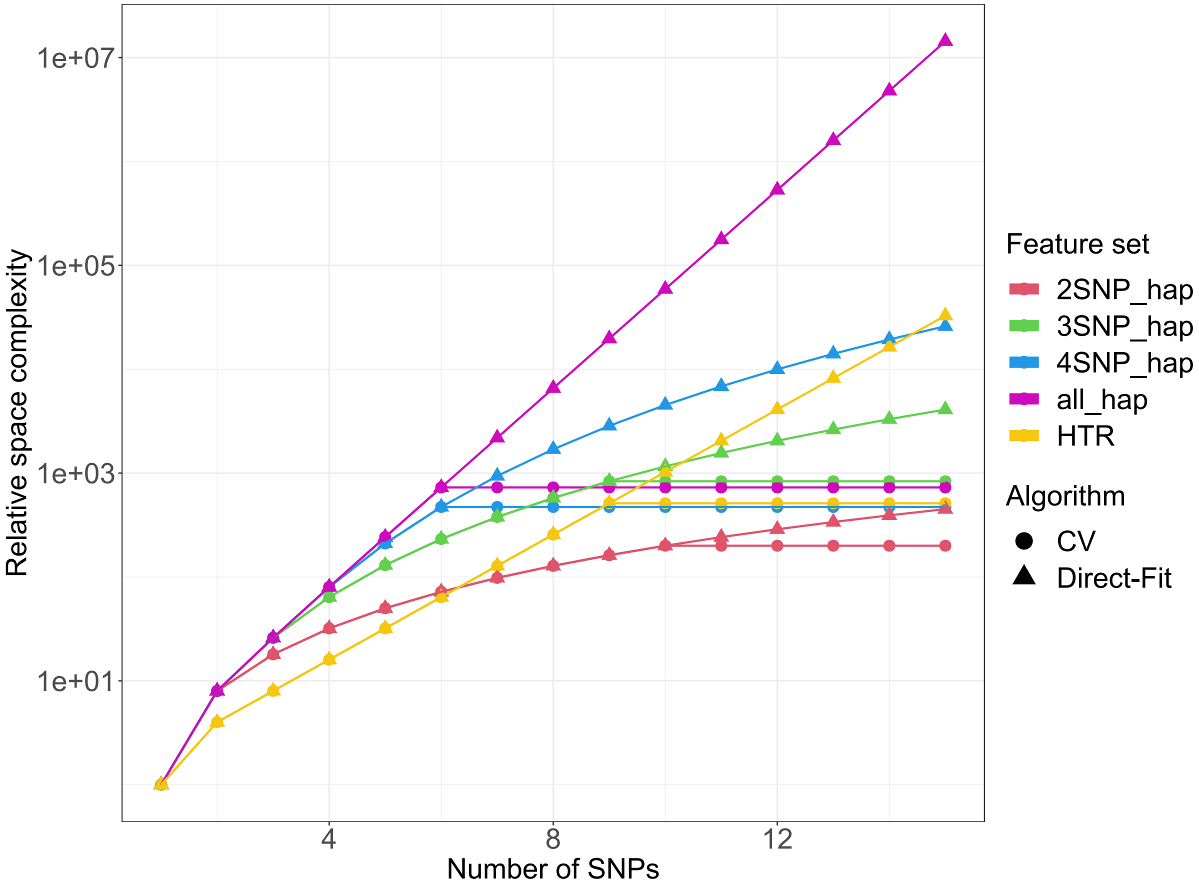


Figure S1: Comparison of space complexity for Algorithm Direct-Fit and CV (the combination of 'Two-stage CV' and 'Cumulative HTRX' for CV) on different feature sets for both linear and logistic regression models. Feature set specifies the maximum number of features (of 6) that can interact, from 'SNP'= 1, to 'all_hap'= 6, while 'HTR' uses templates that interact in all features with no 'X' in the template. The space complexity for linear and logistic regression models is approximately proportional to the number of the input features, which is suggested to be below 1000 for 'CV'. Algorithm 'Direct-Fit' and 'CV' (using 'Two-stage CV') begin with the same number of features when the number of SNPs is smaller than 6. When the number of SNPs increases, the number of features increases exponentially, and 'CV' takes advantage of the Algorithm 'Cumulative HTRX for CV' to reduce the space complexity significantly. When using haplotypes with at most 2 or 3 SNPs ('2SNP_hap' and '3SNP_hap'), or all the SNPs ('HTR'), 'Two-stage CV' allows the region spanning more SNPs while keeping the relative space complexity below 1000.

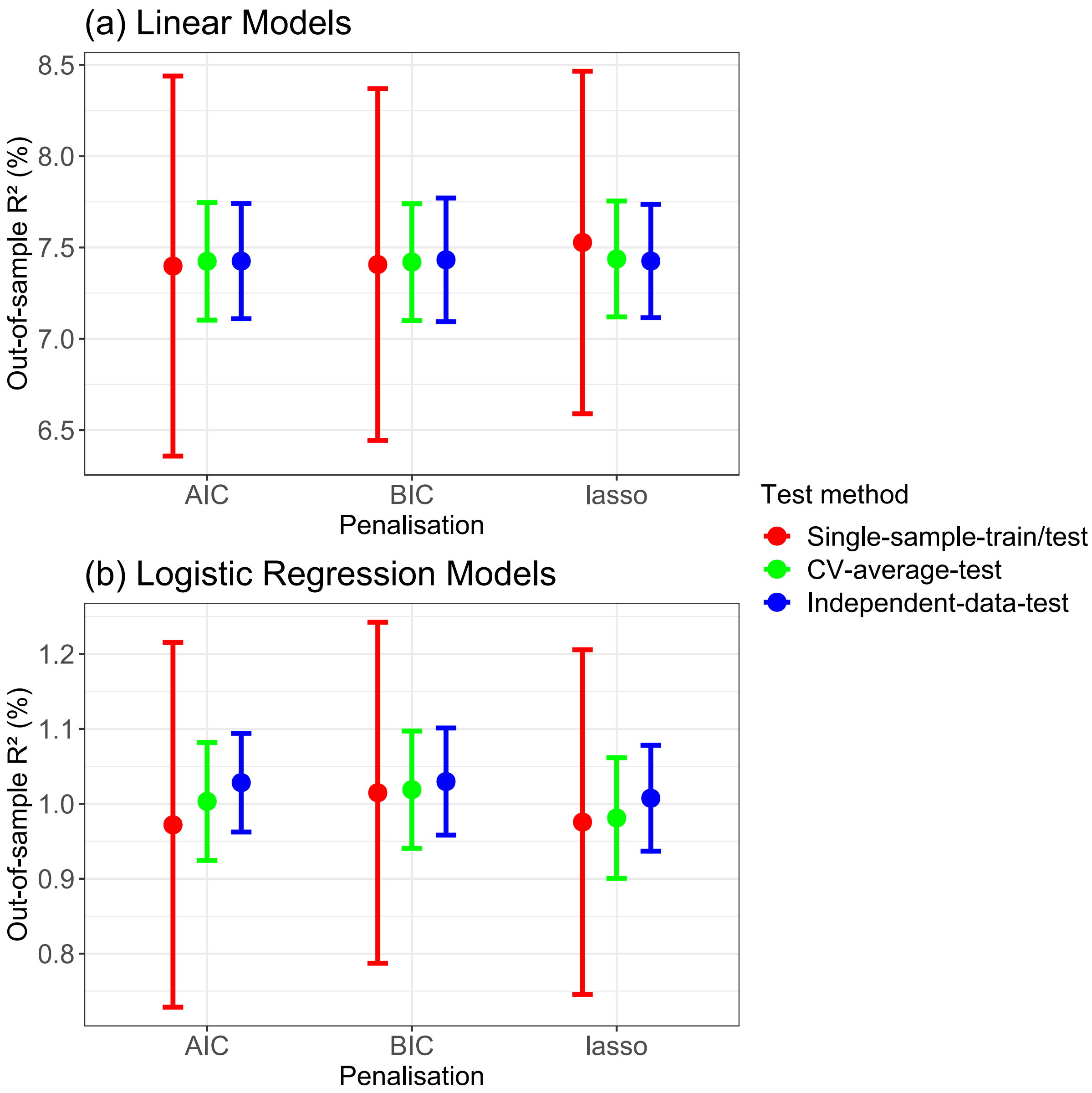


Figure S2: Comparison of out-of-sample variance explained from different test methods. The round dots represent the average of 250 out-of-sample $R^2$ from simulation, and the error bar shows the distance of one standard deviation from the average of 250 simulated out-of-sample $R^2$. 'Single-sample-train/test', 'CV-average-test' and 'Independent-data-test' represent different test methods specified in Supplementary Methods 1.2 (1), (2) and (3), respectively. (2) is our recommended CV approach for real data (Algorithm 2). (1) is a traditional test-train split of that data, and (3) requires additional out-of-sample data.
